# A Machine Learning Model of Perturb-Seq Data for Use in Space Flight Gene Expression Profile Analysis

**DOI:** 10.1101/2024.11.28.625741

**Authors:** Liam Johnson, James Casaletto, Sylvain Costes, Caitlin Proctor, Lauren Sanders

## Abstract

The genetic perturbations caused by spaceflight on biological systems tend to have a system-wide effect which is often difficult to deconvolute into individual signals with specific points of origin. Single cell multi-omic data can provide a profile of the perturbational effects but does not necessarily indicate the initial point of interference within a network. The objective of this project is to take advantage of large scale and genome-wide perturbational or Perturb-Seq datasets by using them to pre-train a generalist machine learning model that is capable of predicting the effects of unseen perturbations in new data. Perturb-Seq datasets are large libraries of single cell RNA sequencing data collected from CRISPR knock out screens in cell culture. The advent of generative machine learning algorithms, particularly transformers, make it an ideal time to re-assess large scale data libraries in order to grasp cell and even organism-wide genomic expression motifs. By tailoring an algorithm to learn the downstream effects of the genetic perturbations, we present a pre-trained generalist model capable of predicting the effects of multiple perturbations in combination, locating points of origin for perturbation in new datasets, predicting the effects of known perturbations in new datasets, and annotation of large-scale network motifs. We demonstrate the utility of this model by identifying key perturbational signatures in RNA sequencing data from spaceflown biological samples from the NASA Open Science Data Repository.

## INTRODUCTION

Recently, efforts to probe the full extent of the influence spaceflight has on gene expression have begun to reveal the gravity of the effects. The NASA Twins Study, a landmark endeavor to provide a comprehensive and multi-omic map of spaceflight effects on the human body, found that after a 6 month stay in space nearly 1,500 genes were differentially expressed between astronaut Scott Kelly and his Earth-bound twin Mark Kelly (Garrett-Bakelman et al., 2019). Additionally, over 100 of these differentially expressed genes (DEGs) did not return to nominal levels post flight. Increasing the in-flight time from 6 to 12 months induced a sixfold leap in the number of DEGs to over 8,500: close to half the protein coding genes in the human genome. The subset of genes which did not revert to normal upon return also increased to over 800. The findings of the Twins Study paint a sobering picture of the long-term effects of spaceflight on the human body and highlight the critical importance of understanding exactly how and why our genes react in response to an extraterrestrial environment.

Out of the more than 8,500 DEGs documented by the Twins Study, it is probable that only a much smaller subset of them were being directly affected by the environment. While bulk and single-cell RNA sequencing both provide a broad and relatively unbiased snapshot of transcription levels, a captured shift in the expression of any given gene in response to spaceflight is only correlative, not causal. Despite being comprised of modular signal motifs which are largely transparent when examined in isolation, gene expression networks are highly interdependent. When the expression of one gene is altered due to an external perturbation, any genes further down may be shifted in a cascade effect. To further complicate matters, the sensitivity of the local motif may result in a situation where the expression of the directly affected gene is shifted only slightly-perhaps not enough to be considered significant-while the downstream genes caught in the cascade cross the threshold of significance easily. Add to these considerations the fact that multiple gene networks can intersect, and endeavoring to derive which genes are being directly perturbed by the environment from expression data alone becomes a task for which classical computation is poorly suited. Machine learning (ML), on the other hand, provides a more promising path forward (Xu & Jackson, 2019). However, ML algorithms require a great volume of data on which to train, which can be problematic when the data is sourced from experiments conducted in space.

Biological research in space is subject to many unique constraints related largely to the logistical limitations of conducting physical experiments in orbit or beyond, resulting in datasets with small sample size. In omics research especially, the feature count can be orders of magnitude larger than the sample count. Transfer learning is a technique to train on large amounts of data from a related domain, thereby creating a pre-trained or generalized model that is already primed to analyze the limited set of spaceflight data. Implementing a pre-trained model capable of making predictions on gene expression data, however, involves a unique challenge: finding a compatible dataset that is large enough to encapsulate complex interactions within gene expression networks.

The Perturb-Seq technique was first established in 2016 as an improved method of genetic screening in mammalian cells with the specific goal of creating high throughput assays of complex phenotypes in the form of transcriptional profiles (Dixit et al., 2016). The principle behind the technique was derived largely from the Synthetic Genetic Array, which served a similar purpose in yeast, but was limited to macroscopic observation rather than molecular data collection (Tong et al., 2001). Many different Perturb-Seq datasets have since been created, primarily in human and murine cell lines. The basic principle of Perturb-Seq involves creating a knockout using a guided molecular technique, usually a form of CRISPR delivered via lentiviral pooling, to knock out a set of genes in a pool of cells. After a sufficient length of time to allow transcriptional responses to occur, the cells are then collected and processed via single cell RNA sequencing (scRNA-seq).

Perturb-Seq datasets, especially ones which knock out and record the responses of thousands of genes, tend to have extremely high dimensionality. This can make them difficult to analyze though traditional computational models. Several variations of Principal Component Analysis (PCA) can be used to cluster the datasets and reduce the number of dimensions, but the resultant clusters often lack any visibly meaningful spatial distinction due to the original dimensionality and volume of the data. Consequently the concept of training machine learning algorithms on Perturb-Seq data is not a novel one (Ji et al., 2021).

There are a number of systems created specifically for this purpose such as Deep Velo, and Cell Oracle, as well as many examples of non-specialized ML architectures being applied to great effect (Kamimoto et al., 2023),(Chen et al., 2022). However, many of these models are created or configured specifically to focus on temporal patterns in perturbational data in order to gain greater understanding of the metabolic structures which produced the data, rather than to predict sources of an effect in unrelated data (Tegnér et al., 2003). Consequently, constructing and optimizing a model from the ground up was determined to be a more prudent course of action than attempting to adapt one which had been tuned to a different purpose.

Here we report the results of pre-training three types of generalized machine learning model on a Perturb-Seq dataset constructed from a human cell line, then used to make predictions on smaller, spaceflight data in order to identify the upstream sites of perturbation responsible for the widespread transcription alterations that occur due to spaceflight. Two of the models tested in particular display a promising capacity for making informative predictions despite the various differences in quantity, quality, and nature which separate the original training data from the samples used for prediction.

## METHODS

We train a series of machine learning models to predict which gene is knocked out in each cell of a Perturb-Seq dataset (Replogle et al, 2022); or to predict which biological function appears to be most strongly perturbed in that cell via clustering. After training the model on a large number of cells, we then predict either the specific gene perturbation, or the strongest biological perturbation, within space biology RNA-seq samples from NASA’s Open Science Data Repository (OSDR). Although the training data for the model must be single cell data by necessity, prediction is possible for both single cell and bulk RNA-seq data by treating the bulk sample as a single datapoint. All programming was done in Python as the language is ideal for both bioinformatics and high-level machine learning.

### Genome-Wide Perturb-Seq

The Genome-Wide Perturb-Seq (GWPS) datasets were accessed at https://gwps.wi.mit.edu/. This database includes 3 scRNAseq datasets: e genome-wide knockouts in the K562 chronic myeloid leukemia cell line (K562GW); a subset of the K562W dataset containing only the perturbations which had resulted in transcriptional profiles with significant responses, termed the K562 Essential Genes dataset (K562E); and a third dataset of gene knock-outs in the non-cancerous retinal pigment epithelial cell line RPE1. Notably, the K562E and RPE1 datasets are of a similar size and design, making them suitable for direct performance comparison (Table 1).

**Table 1.** Overview of the Genome-Wide Perturb-Seq scRNA-Seq data. Cell Count: the number of cells sequenced from each cell line. Filtered Cell Count: cell count after filtering using Scanpy to remove cells with no genes knocked out or any other quality defects. Perturbations: the number of genes knocked out in each cell line. Transcriptional profile: the number of genes with expression quantified via scRNAseq in each cell line.

| Overview of the Genome-Wide Perturb-Seq Library |  |  |  |  |
| --- | --- | --- | --- | --- |
| Dataset | Cell Count | Filtered Cell Count | Perturbations | Transcriptional Profile |
| K562WG | 1,989,578 | N/A | 9,854 | 8,248 |
| K562E | 310,385 | 299,694 | 2,058 | 8,563 |
| RPE1 | 247,914 | 236,429 | 2,392 | 8,749 |

The K562E and RPE1 datasets were chosen as the input to the machine learning models due to the similarity of their size and knockout coverage. The complete distributions of perturbed genes and their cell counts are plotted in Figure 1C.

**Figure 1.**
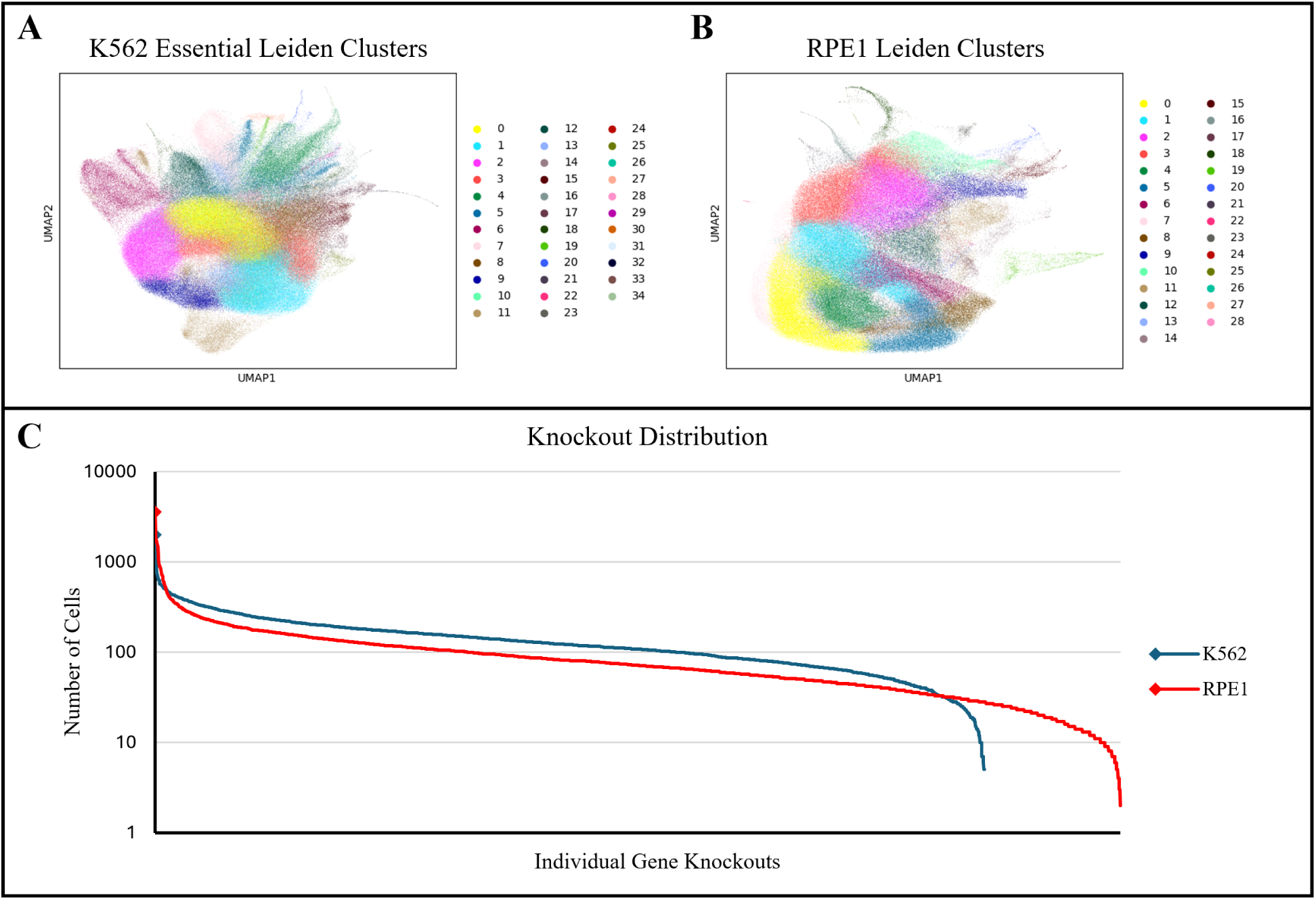
GWPS clusters and knockout distributions. (**A & B**) The results of Leiden based Uniform Manifold Approximation and Projection (UMAP) clustering for K562E and RPE1 Datasets, which resulted in 35 and 29 clusters respectively. (**C**) A graph of the cell counts for each of the over 2,000 knockouts present in the two datasets. The distribution is highly biased and is shown on a logarithmic scale in order to better visualize the inconsistency.

### Clustering

Both the K562E and RPE1 datasets were first filtered as a preliminary step in order to remove cells which did not receive a knockout, which removed approximately 10,000 cells from each. This was done to prevent the formation of an un-perturbed category in the gene list from influencing both cluster formation and direct knockout prediction. The datasets were then clustered using the Scanpy python package (Wolf et al., 2018). The clustering algorithm used for all analysis was a Leiden-driven Uniform Manifold Approximation and Projection (UMAP). The clustering results are shown in Figure 1A-B.

### Machine Learning

Our predictive model needed to be created from an ML algorithm that permitted multiple classes for classification and had a flexible architecture. In total, three algorithms were adapted for trials: a Random Forest distribution from Scikit-Learn, a Multi-Layer Perceptron distribution also from Scikit-Learn, and a Convolutional Neural Network distribution from Keras by way of TensorFlow (TF).

### Data Labeling

Creating predictive models from the GWPS data requires multi-class algorithms that can be configured to sort the data into categories or classes based on labels. We trained classifiers using the knocked-out genes as labels (hereafter referred to as “gene labels”). Because this creates a very high number of classes (∼2000-2300 knocked out genes), model accuracy suffers. In order to address this, we also clustered the scRNAseq data, resulting in approximately 30 clusters per dataset, and annotated each cluster by the biological functions enriched in the top knocked-out genes in that cluster as follows.

The top 50 most common knockouts in each cluster were collected and used as the input for gene set enrichment analysis (GSEA) in order to determine the most enriched biological functions perturbed within that cluster (Culhane et al., 2012). GSEA gene sets Gene Ontology (GO), GO biological process (GO:BP), GO cellular component (GO:CC), GO molecular function (GO:MF), and Human Phenotype Ontology (HPO). The significance threshold (FDR q-value) was set to 0.05, and the maximum number of reported overlaps was restricted to the top 10. The outputs were then compiled and assessed to create an annotation of the biological function for each cluster (Supplementary Tables 1 and 2). These annotations are hereafter referred to as “cluster labels”.

Additionally, some ML techniques, such as Convolutional Neural Nets, are capable of utilizing multiple types of labels simultaneously, thereby enabling the model to form associations with both label and cluster simultaneously. In that scenario the model would still predict based on a single label type, but the association gained through training with other labels would augment the prediction accuracy.

### Scikit-Learn Random Forest

We implemented the Scikit-Learn distribution of the Random Forest algorithm, primarily for the purpose of establishing a performance baseline for the other models. The Random Forest model was used as a training benchmark and was configured with 100 decision trees, which then vote on a single category in which to classify the output. The model was trained on both the K562 Essential and RPE1 datasets using both types of labels and a 30-70 test-train data ratio, so that the performance with both label types could be compared for each for a total of 4 trained models.

Unfortunately, the gene knockout label models originally did not converge even after training for several days. However, it is possible to set a depth limit for the trees in order to force the model to converge at the cost of some performance, so a depth limit of 10 was set for the gene label models only.

### Scikit-Learn Multi-Layer Perceptron

We implemented the Scikit-Learn distribution of the Multi-Layer Perceptron (SKL-MLP), then conducted Hyperparameter Optimization using the following hyperparameters: epoch number, batch size, alpha value, learning rate, and the hidden layer configuration. Of these, the first three were scalar settings, and their impact on performance can be seen as charts in Figures 2 and 3 along with learning rate type.

**Figure 2.**
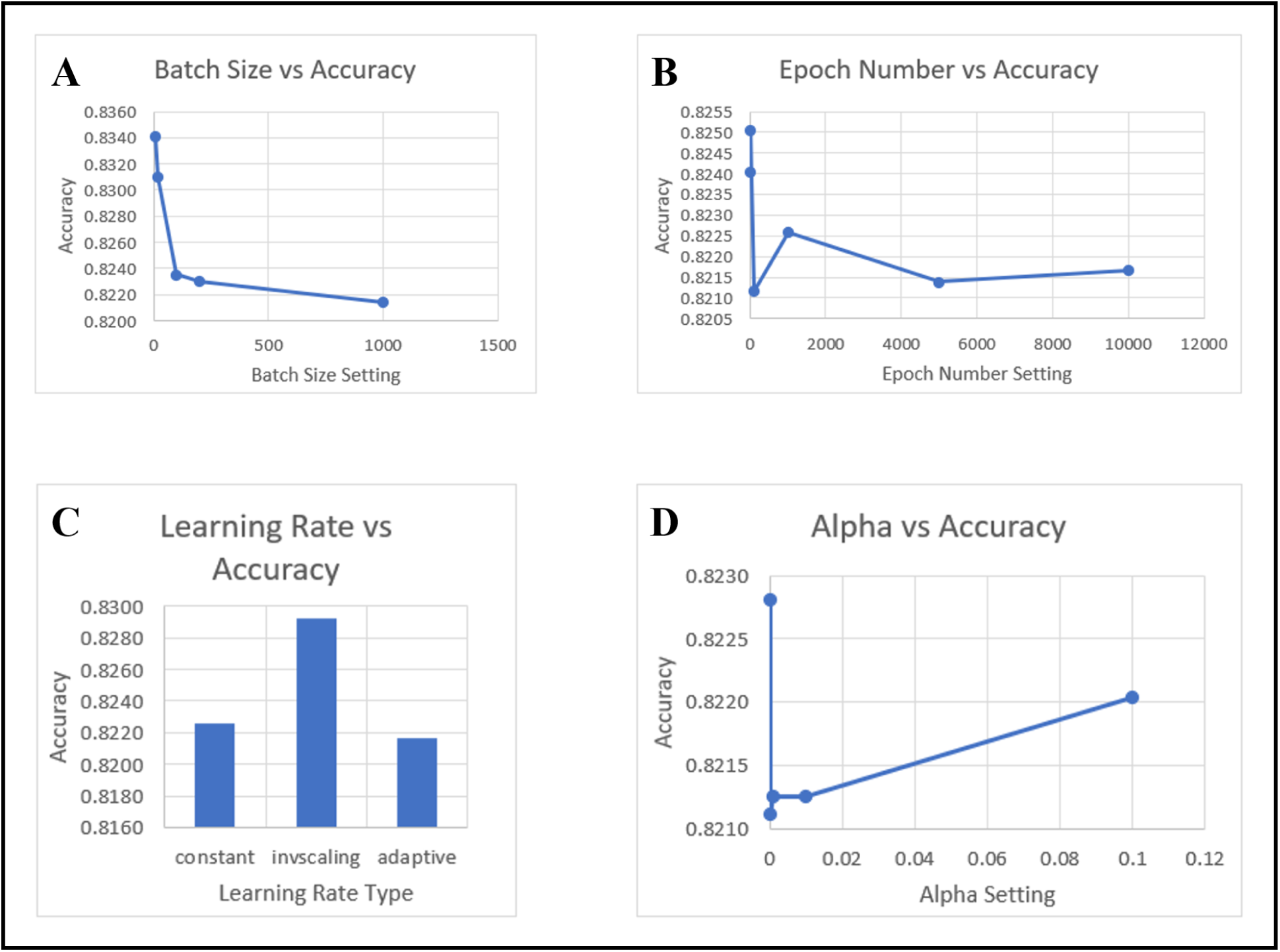
Scikit-Learn Multi-Layer Perceptron hyperparameter accuracy optimization trials. (**A**) The effect of modulating batch size on testing accuracy. (**B**) The effect of the number of training epochs on testing accuracy. (**C**) The effect of different learning rate functions on testing accuracy. (**D**) The effect of modulating alpha value on testing accuracy.

**Figure 3.**
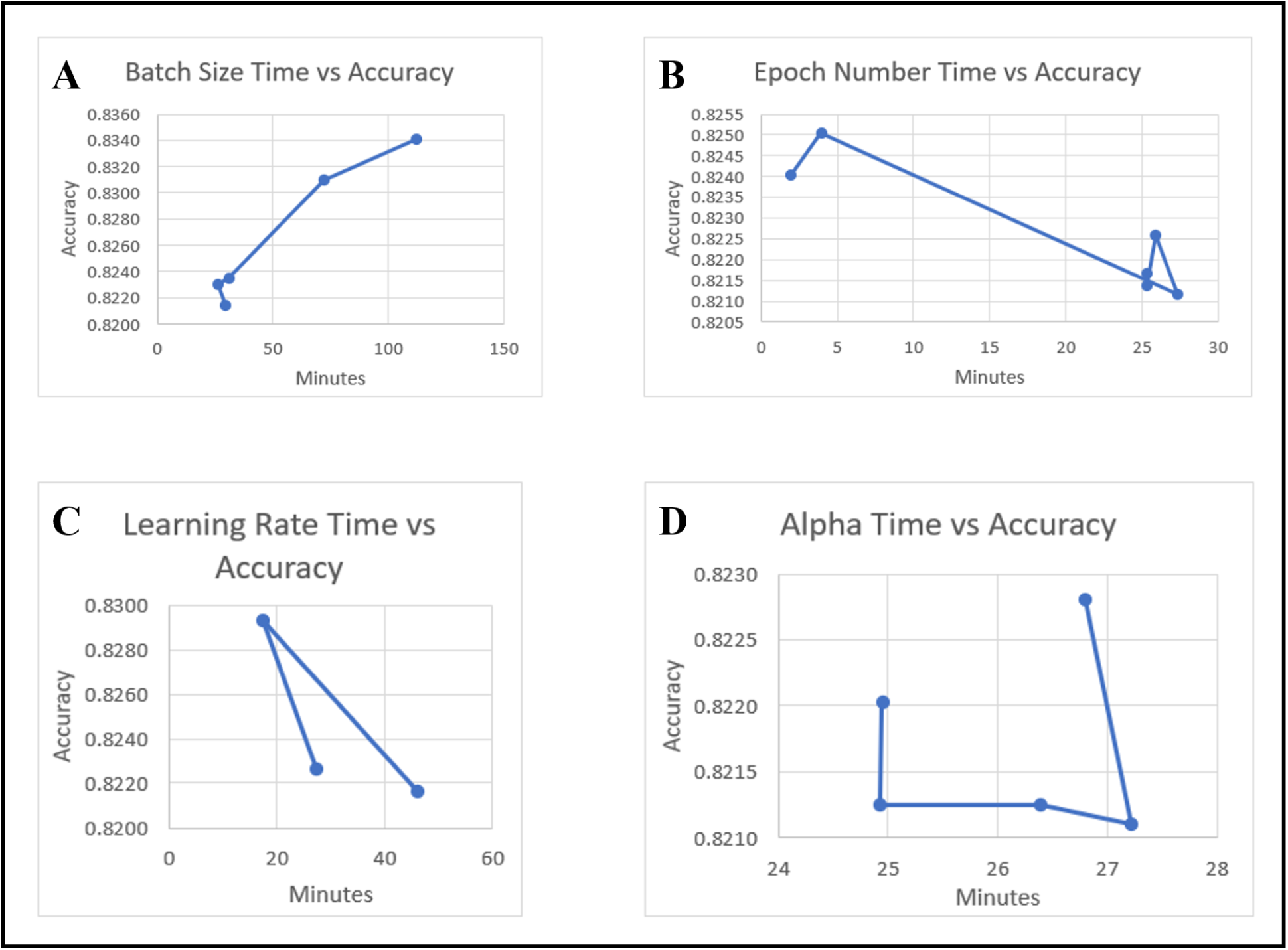
Scikit-Learn Multi-Layer Perceptron hyperparameter accuracy optimization effects on training time. (**A**) The effect of modulating batch size on both testing accuracy and training time. (**B**) The effect of the number of training epochs on both testing accuracy and training time. (**C**) The effect of different learning rate functions on both testing accuracy and training time. (**D**) The effect of modulating alpha value on both testing accuracy and training time.

Tuning the hidden layer configuration is in principle more complicated, as doing so involves adjusting both the number of layers, and the number of nodes within a given layer. The baseline configuration for this setting was chosen by doubling and halving the default value, which was a single layer of 100 nodes. The 200 node version had improved accuracy over the default, but no further refinement to this setting was made. All other configurations tested either performed worse in terms of accuracy, or performed similarly but at the cost of a greatly increased training time. The inclusion of additional hidden layers, whether larger, smaller, or of equal size, also reduced performance without exception. The settings of the final MLP model configuration are in Table 2.

**Table 2.** Scikit-Learn Multi-Layer Perceptron final model structure and configuration.

|  |  |
| --- | --- |
|  | **Input Layer** |
|  | **Hidden Layer:**<br>nodes = 200 |
|  | **Output layer:**<br>nodes = number of classes |
|  | **Model Compilation Settings:**<br>training epochs = 10, optimizer = 'solver', batch size = 10, alpha = 0.1, learning rate = 'invscaling', initial learning rate = 0.001, batch size = 10 |

### TensorFlow-Keras Convolution Neural Network

We implemented a Convolutional Neural Network (CNN) via the TensorFlow application programming interface (API) distribution of the Keras CNN in several configurations, then conducted Hyperparameter Optimization. In addition to the cluster and gene label model variants, we configured the TF-Keras CNN to accept multiple label types simultaneously for the same data, forming a composite model. By training on both cluster and gene knockout labels, we used the associations from both to improve the accuracy of the cluster labels which were used for the output classification.

We performed hyperparameter tuning to generate the optimum CNN configuration, shown in Table 3. Graphs of model behavior during hyperparameter tuning can be found in Figures 4, 5, and 6.

**Figure 4.**
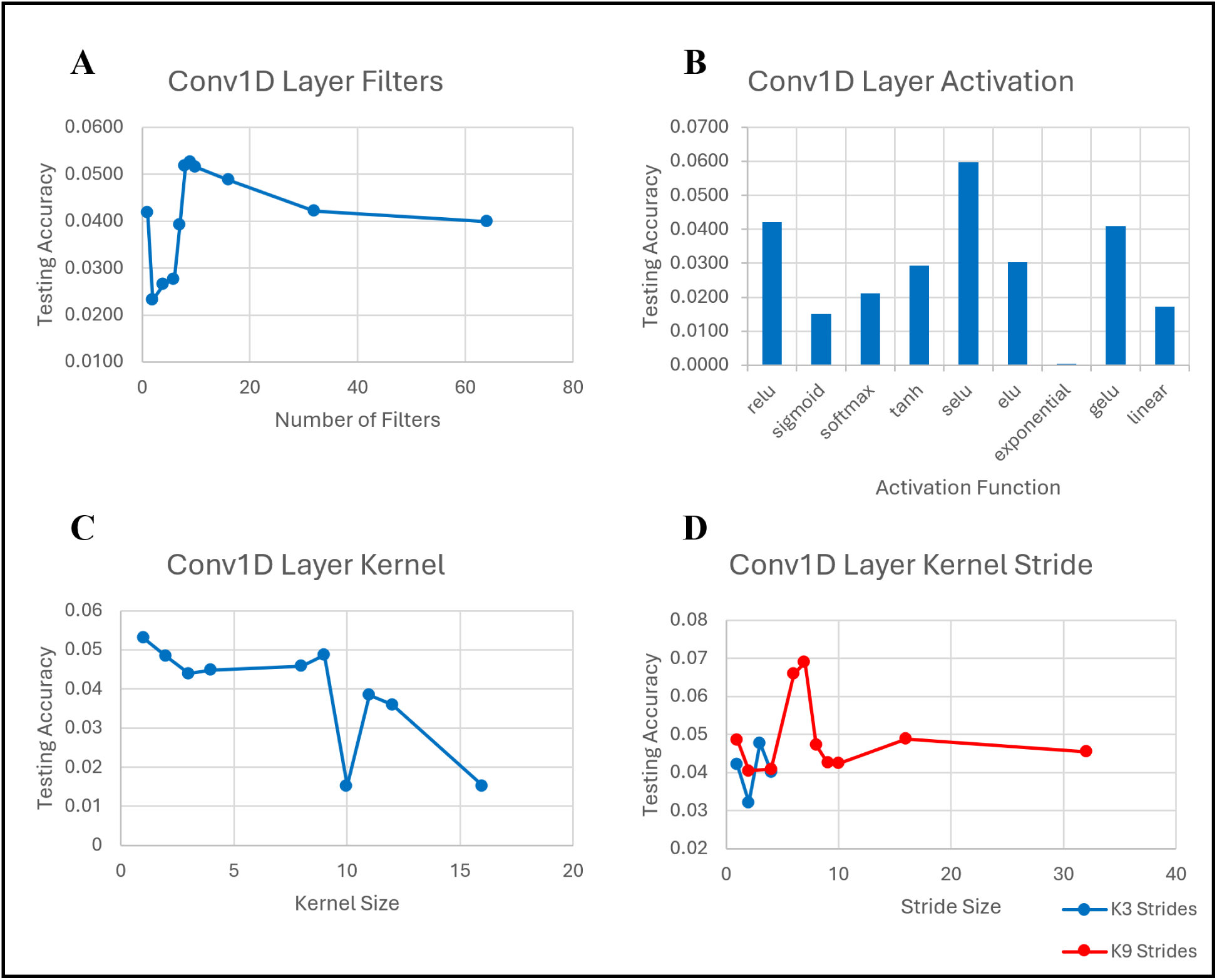
TF-Keras Convolutional Neural Network 1D convolutional layer hyperparameter optimization trials. (**A**) The effect of modulating the number of filters on testing accuracy. (**B**) The effect of different activation functions on testing accuracy. (**C**) The effect of modulating the kernel size on testing accuracy. (**D**) The effect of modulating the width of the kernel stride on testing accuracy. This effect is directly linked to the kernel size, so the behavior for two different kernel sizes is shown.

**Figure 5.**
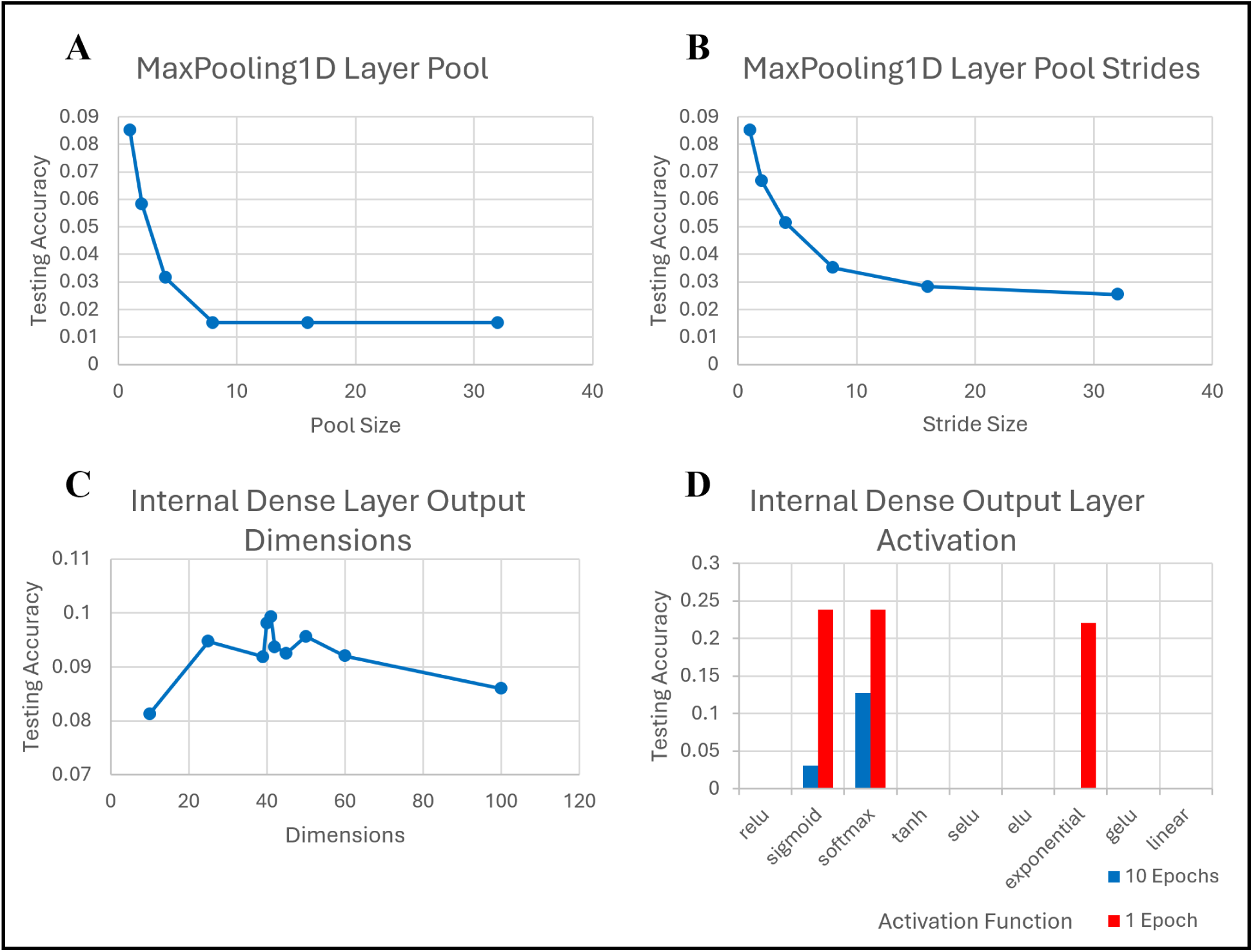
TF-Keras Convolutional Neural Network pooling, dense, and output layer hyperparameter optimization trials. (**A**) The effect of modulating the pool size on testing accuracy. (**B**) The effect of modulating the width of the pool stride on testing accuracy. This effect is directly linked to the pool size, but as the former behaves predictably multiple plots are not required. (**C**) The effect of modulating the number of nodes in the dense layer on testing accuracy. (**D**) The effect of different activation functions on testing accuracy. Sigmoid and Softmax behave differently for probability-based prediction, so the similarity of their performance grants some flexibility in the configuration of a prediction system.

**Figure 6.**
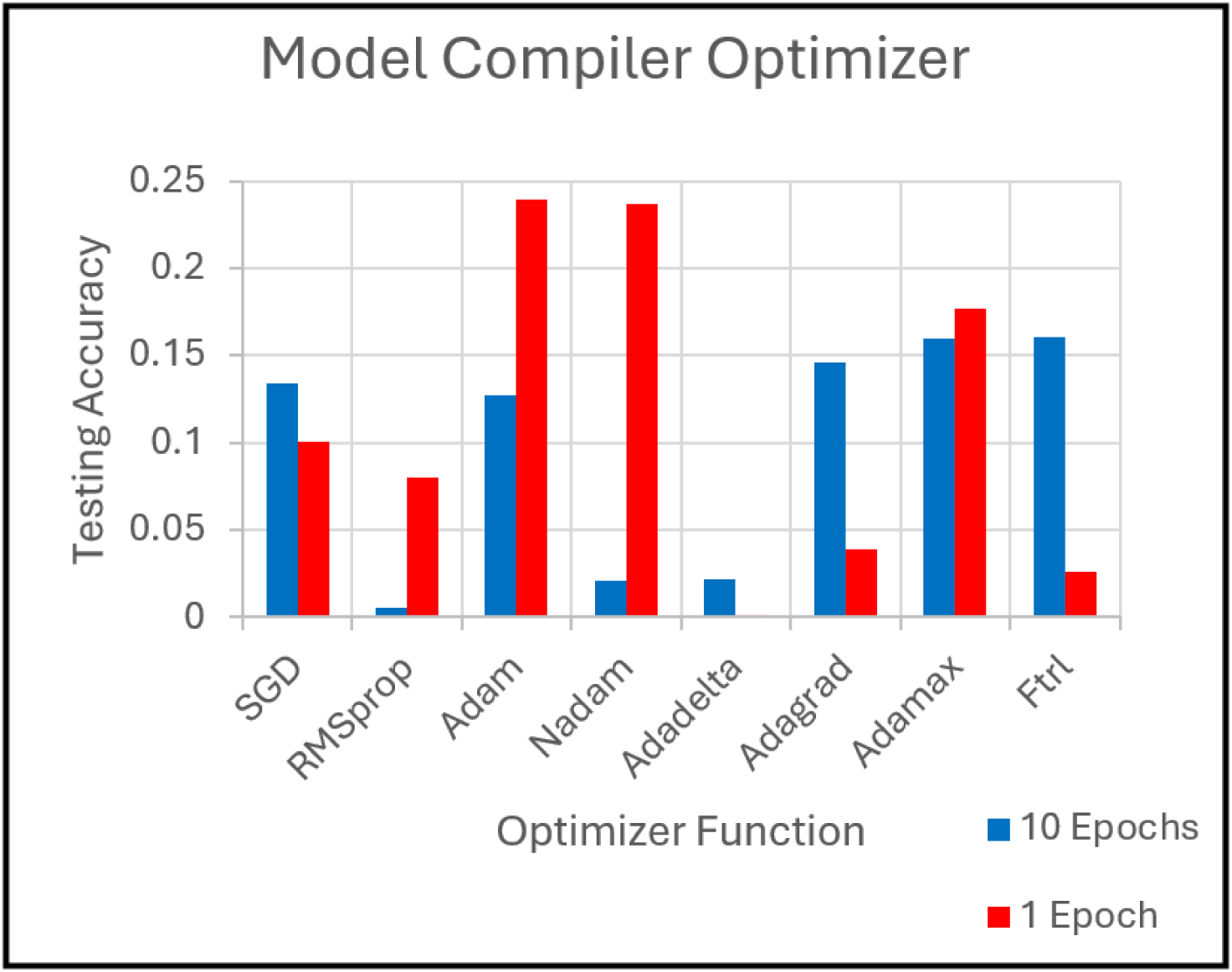
TF-Keras Convolutional Neural Network model compiler hyperparameter optimization trials. The model compiler is not a specific layer, but adjusting it still alters the model performance significantly.

**Table 3.** TF-Keras Convolutional Neural Network final model structure and configuration.

|  |  |
| --- | --- |
|  | **1D Convolutional Layer:**<br>filters = 16, kernel size = 9, strides = 7, activation = 'selu', input shape = data dimensions |
|  | **Flatten Layer** |
|  | **Dense Layer:**<br>units = number of classes, activation = 'sigmoid' |
|  | **Model Compiler:**<br>training epochs = 1, optimizer = 'adam', loss = 'sparse categorical crossentropy' |

### Space Biology Data

Data from NASA’s Open Science Data Repository (OSDR) was used to validate the ability of the MLP models to make new predictions on spaceflight data. Four bulk RNA-seq samples were downloaded from OSD-91, a simulated microgravity experiment conducted in the human lymphoblast cell line TK6. Prediction required the OSD-91 gene expression data to match the gene expression data on which the model was trained. The OSD-91 data had transcriptional profiles that included roughly 16,000 genes. Despite being half the size the two Perturb-Seq transcriptional profiles still included many genes that were not part of the OSD-91 coverage: specifically, 659 genes in the RPE1 dataset, and 462 genes in the K56 dataset. In order to rectify the dimensional disparity, the OSD-91 data was first sheared of any genes not present in the model data. Then, the genes unique to the model data were added to the OSD-91 transcriptional profile, and their differential expression values set to zero in order to ensure that they would not be introducing a false signal. As the K562 and RPE1 models had slightly different dimensions, the process was repeated for both. Cluster and gene predictions for the 4 OSD-91 samples were then made using the models from both the K562 and RPE1 cell types.

## RESULTS

### Perturb-Seq dataset quality can be normalized, compared, and evaluated for applicability of use as training data

In order to assess the overall distribution of gene knock-outs in each dataset, we performed an exploratory data analysis prior to model training. Ideally, a training dataset would include scRNAseq data from the same number of cells from each gene knock-out. However, due to the nature of the Perturb-Seq assay, the distribution of each gene knocked-out across the dataset is extremely variable.

Therefore, here we establish a method of evaluating single multiplicity of infection (MOI) Perturb-Seq data quality based on cluster formation and composition that both enables the comparison of different datasets and provides a means of assessing the impact data quality may have on model performance. We use cluster formation as a proxy for broad transcriptional effects, relying on the assumption that a computationally derived cluster should be mainly composed of cells with the same gene knocked-out; or at least composed of cells with genes from the same functional network knocked-out.

First, the number of cells in each cluster is calculated by referencing the parent cluster for each cell in the dataset. Then the counts are normalized by dividing them by the total number of cells in the dataset. The process is then repeated to determine the number of unique knockouts that occur in each cluster, which are normalized through division by the knockout coverage (total number of unique knockouts). It is important to note that cluster size can inform this to some extent, particularly in smaller clusters as they may be composed of fewer cells than the number of possible knockouts. Then, the internal uniformity of knockouts for each cluster can be determined by dividing the normalized size by the normalized knockout distribution. The outputs of this process for the K562E and RPE1 datasets can be seen in parts A, B, and C respectively of Figure 7. The value for internal cluster uniformity can range from 0-1, with 1 being a completely homogeneous cluster where only 1 gene is knocked out in all cells in that cluster. Averaging the values across all clusters then results in a single quality score for the dataset.

**Figure 7.**
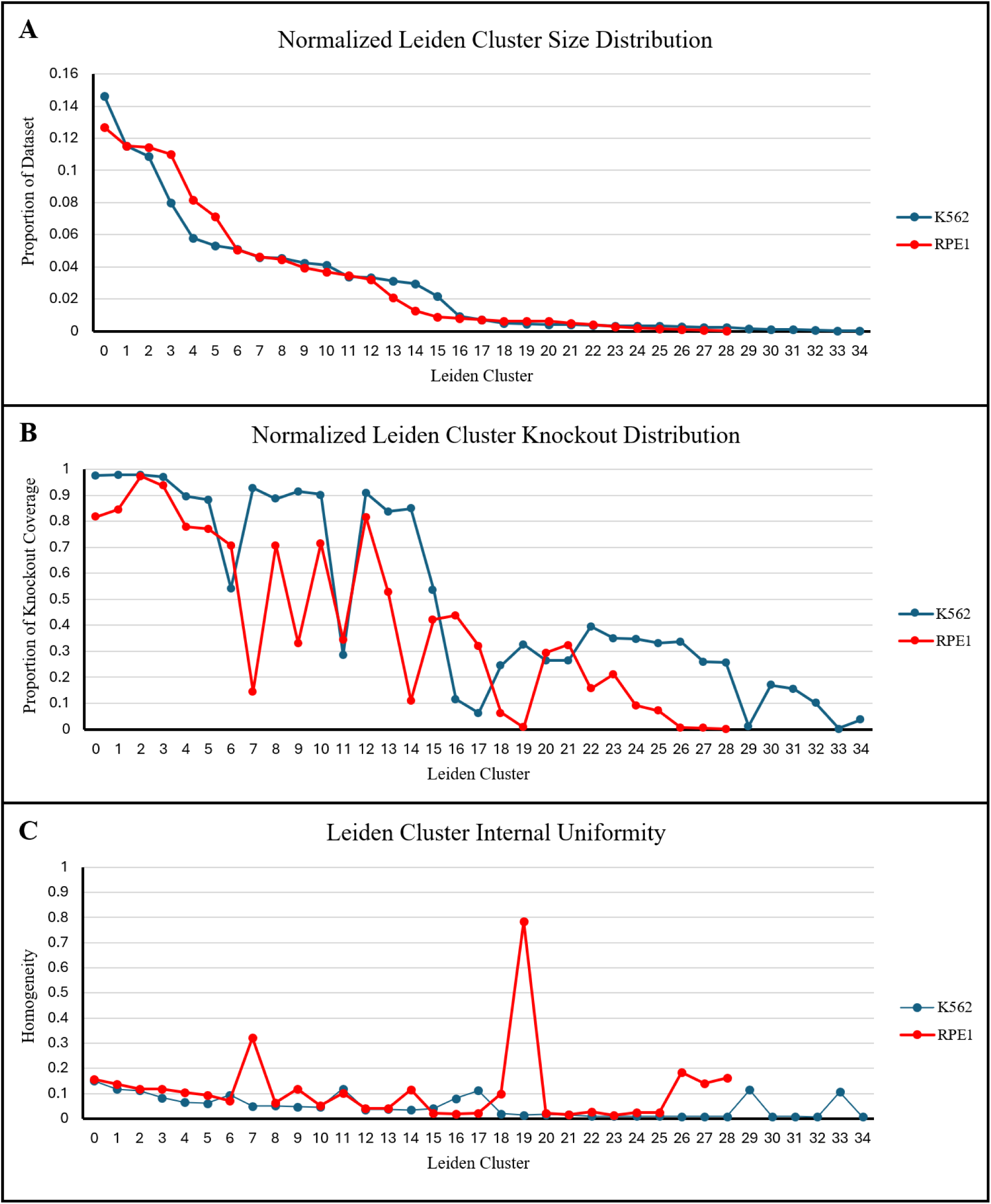
Quality assessment of the K562E and RPE1 datasets. (**A**) The graph of the cell counts for each cluster in the two datasets, normalized by dataset size and ordered by cluster size. (**B**) The graph of the proportion of possible knockouts that occur in each cluster. (**C**) The graph of the level of internal uniformity (homogeneity) of genes knocked out within each cluster, the average of which is the quality score.

By this metric, the K562-derived Essential Genes dataset had a quality score of 0.049 while the RPE1-derived dataset had a score of 0.111, implying that the transcriptional effects caused by knocking out genes in the RPE1 cell line resulted in a dataset that could be grouped into distinct clusters more effectively.

Although a quality score based on the internal uniformity of the clusters does not simultaneously encompass other difficult to capture aspects such as cross-cluster redundancy, it provides a stable metric to directly assess the datasets beyond the distribution of knockout cell counts.

The gene set enrichment analysis and resultant cluster annotations for each dataset mirrors the quality assessments as well, with the K562E clusters tending to have much greater repetition of function between them, while we would instead expect each cluster to be functionally distinct given that clusters should be capturing the transcriptional effects from different knocked-out genes. The RPE1 clusters on the other hand are noticeably more distinct from each other, and have a far greater coverage of gene sets linked to distinct processes and human health implications. The annotations are available for comparison in Supplementary Tables 3 and 4 respectively. While the ubiquitous presence of mitotic cell cycle regulation and processes in the annotations of both datasets is to a large extent unavoidable due to a combination of immortal cell line behavior and the tendency of scRNA-seq to favor cell-cycle transcripts (Barron & Li, 2016), their presence and is visibly lower in the non-cancerous RPE1 cell line in comparison to the cancerous K562E line. Taken together, the quality and annotation behavior of the two comparable datasets in conjunction with known attributes such as consistency of ploidy level and stability of expression landscape suggests that the K562 cell line may be unsuited to Perturb-Seq applications (Lombardi et al., 2022). However, we perform model training and downstream analysis on both cell lines, to enable comparison.

### Multi-Layer Perceptron provides highest accuracy and reliability for predicting gene perturbations

The SKL Random Forest, SKL-MLP, and TF-Keras CNN models were all compiled and trained on both the K562E and RPE1-derived datasets. Different variations of the models used either the knocked out genes or the associated clusters as labels, or both simultaneously in the case of the composite variant of the TF-Keras CNN. This resulted in a total of 8 generalized models for comparison.

Universally, the ML models trained on RPE1 data exhibited a higher performance than those trained on K562 data, regardless of architecture and label type as seen in Table 4. Notably, this is consistent with the quality scores for the respective datasets. For both datasets, the SKL-MLP cluster label models were the most accurate out of the three ML algorithm types implemented. When gene knockouts were used as labels, the SKL-MLP performed well, but was superseded by the TF-Keras CNN.

**Table 4.** Testing accuracies of all machine learning models. Each of the three machine learning algorithms was trained on each of the two datasets using each type of compatible label. All reported testing accuracies are an average of the testing accuracies for each class within the model.

| **Testing Accuracy of K562 Models** |  |  |  |
| --- | --- | --- | --- |
| **Algorithm** | **Labels** |  |  |
|  | **Cluster** | **Gene Knockout** | **Composite** |
| SKL Random Forest | 61.7% | 2.4% | N/A |
| SKL-MLP | 83.3% | 6.6% | N/A |
| TF-Keras CNN | 79.2% | 14.4% | 73.0% |
| **Testing Accuracy of RPE1 Models** |  |  |  |

| Algorithm | Labels |  |  |
| --- | --- | --- | --- |
|  | Cluster | Gene Knockout | Composite |
| SKL Random Forest | 70.2% | 3.8% | N/A |
| SKL-MLP | 87.7% | 9.6% | N/A |
| TF-Keras CNN | 83.0% | 23.9% | 83.6% |

While the testing accuracies of the models trained using gene knockouts as labels were generally much lower than the models trained using clusters as labels, it is insufficient to use the scores as a direct comparison, primarily due to the number of classes (Choromanska et al., 2013). The SKL-MLP RPE1 gene model, for example, achieved a testing accuracy of 9.6% which seems low, but is in fact a roughly 240-fold improvement over the null model accuracy of 0.04 % (1/2,392). By contrast the cluster label version, despite achieving a testing accuracy of 87.7%, is only a 24-fold improvement over the 3.6% (1/28) accuracy of its null model. The difference in performance of the TF-Keras CNN is even more extreme. Although it attained similar results when the clusters were used as labels, the gene knockout model had a test accuracy of 23.9%: a nearly 600-fold improvement over the null model accuracy. Despite higher accuracy, the TF-Keras CNN models reached an optimal output after only a single training epoch, which is a potential indicator for overfitting. Additionally, the advantage yielded by the composite CNN configuration was relatively insignificant. For these reasons, the SKL-MLP was determined to be the more reliable of the two and so was chosen to be adapted for the prediction framework. The performance values for all models are reported in Table 4.

### Predictions made on simulated space biology datasets

Due to the performance and reliability of the SKL-MLP, we selected this model to predict gene knockouts and functional perturbations in the space biology samples from the OSD-91 dataset (Chowdhury, 2016). This dataset contains four bulk RNA-seq samples (two experimental replicates and two control replicates) from TK6 cells which had been exposed to simulated microgravity for 48 hours in a rotating bioreactor (Chowdhury et al., 2016).

Due to the fact that the OSD-91 samples were perturbed by complex environmental stressors, it is likely that multiple genes are affected directly. Therefore, we report and assess the top 5 gene and cluster predictions.

Then, since the top 5 gene and cluster predictions for each sample were the same, we report the average probability of both labels across all 4 samples (Tables 5 & 6). In order to assess the biological consistency of the gene and cluster predictions, we investigated whether the top 5 predicted genes were among the 50 most frequently knocked out genes in each cluster. This assessment could also potentially reinforce the prediction of the low-accuracy gene label model.

**Table 5.** K562E model predictions on OSD-91 data. Here all prediction metrics and cross-references for the K562E models are reported. Included are the top 5 most probable cluster matches, the top five most probable gene matches, and a check of whether the top gene matches occur in the top 50 most frequent knockouts that the two most probable clusters are comprised of. The latter serves as a cross-reference validation between the two models.

| **K562E SKL-MLP Model Predictions for OSD-91 Samples** |  |  |  |  |  |  |
| --- | --- | --- | --- | --- | --- | --- |
| **Cluster Labels** |  |  | **Gene Knockout Labels** |  | **Rank in Cluster 2** | **Rank in Cluster 4** |
| **Annotation** | **Cluster** | **Avg. Probability** | **Perturbation** | **Avg. Probability** |  |  |
| mitotic cell cycle processes | 2 | 25.18% | *SF3A2* | 1.3% | Not in top 50 | 17<sup>th</sup> |
| multiple stages of mRNA processing, formation and operation of spliceosome | 4 | 18.42% | *NACA* | 0.8% | Not in top 50 | Not in top 50 |
| mitotic cell cycle processes, chromosome organization and segregation | 0 | 17.54% | *DDX46* | 0.7% | Not in top 50 | 43<sup>rd</sup> |
| cell cycle regulation, replisome, replication fork | 18 | 11.53% | *SART1* | 0.6% | Not in top 50 | 16<sup>th</sup> |
| mitotic cell cycle processes, transcription initiation | 3 | 8.36% | *SF3B2* | 0.6% | Not in top 50 | 9<sup>th</sup> |

**Table 6.** RPE1 Model Predictions on OSD-91 Data. Here all prediction metrics and cross-references for the RPE1 models are reported. Included are the top 5 most probable cluster matches, the top five most probable gene matches, and a check of whether the top gene matches occur in the top 50 most frequent knockouts that the two most probable clusters are comprised of.

| RPE1 SKL-MLP Model Predictions for OSD-91 Samples |  |  |  |
| --- | --- | --- | --- |
| Cluster Labels | Gene Knockout Labels | Rank in Cluster 0 | Rank in Cluster 24 |

| Annotation | Cluster | Avg. Probability | Perturbation | Avg. Probability |  |  |
| --- | --- | --- | --- | --- | --- | --- |
| RNA processing and integrator complex, cell cycle – membrane separation, abnormal mitochondrial morphology, mental deterioration | 0 | 81.2% | *PPP6C* | 19.0% | 8<sup>th</sup> | 15<sup>th</sup> |
| mRNA and RNA processing and maturation, histone encoding, extranuclear transport | 24 | 7.3% | *SLC1A5* | 4.1% | 2<sup>nd</sup> | Not in top 50 |
| multivesicular body and vesicle organization and transport, membrane fission | 20 | 5.6% | *MYBL2* | 2.0% | 9<sup>th</sup> | Not in top 50 |
| protein containing complex assembly, abnormal mitochondrial morphology, cytoskeleton organization | 5 | 2.3% | *GAB2* | 1.0% | 11<sup>th</sup> | Not in top 50 |
| supramolecular complex | 1 | 1.5% | *FAM136A* | 0.9% | 20<sup>th</sup> | Not in top 50 |

Table 5 shows the results of the K562-trained model for predictions. The MLP identified Cluster 2 as the highest probability cluster, closely followed by Cluster 4 and Cluster 0. Interestingly, Cluster 4 contains 4 of the top 5 predicted genes within its 50 most frequently knocked out genes, indicating that the biological perturbation of Cluster 4 is more consistent with the gene expression of the OSD-91 samples. This may be due to the lower quality of the K562 Perturb-Seq dataset.

Like many of the clusters in the K562E cell line, the top knockouts in Cluster 2 are enriched for gene sets that drive various functions and stages of the mitotic cell cycle on both a regulatory and biomechanical level. In Cluster 4, the top knockouts are enriched for gene sets that encompass several specific stages of mRNA processing, particularly those involving the formation and operation of the spliceosome. The gene set enrichment analysis results for these clusters are in supplementary table 1.

The original study that analyzed the OSD-91 dataset includes lists of genes found to have differentially hydroxymethylated regions (DHMRs), which are one type of epigenetic modification responsible for differential levels of gene expression (Chowdhury et al., 2016). Thus, one way to validate the performance of our pre-trained models is to compare whether the genes predicted to be knocked-out are functionally consistent with the genes found to have DHMRs in the original study, The K562E gene model predictions do not contain any of the specific genes reported to have the DHMR behavior in the original study. However, the genes reported to have DHMRs broadly included a set of RNA processing and binding factors (Chowdhury et al., 2016). This is consistent with the functions of several of the genes that our pre-trained MLP model predicted to be perturbed in the OSD-91 samples. Of the predicted genes, SF3A2 encodes splicing factor 3a subunit 2, NACA is a nascent polypeptide associated complex subunit, DDX46 is a DEAD-box helicase, SART1 is a spliceosome associated factor as well as a recruiter of U4/U6.U5 tri-snRNP, and SF3B2 is a specific splicing factor (NCBI, 2024).

The predictions made by the models trained on the RPE1 dataset, shown in Table 3.3 were more internally consistent than those of the K562E models. The OSD-91 samples were predicted to belong to Cluster 0 with 81.2% probability, nearly as high as the original training accuracy for the RPE1 MPL cluster labels model (87.7%). Additionally, the predictions of the model trained on gene knockout labels agree strongly with the predictions of the model trained on cluster labels, as each of the top 5 predicted genes is present in the 50 most frequently knocked out genes in Cluster 0. On their own, the sources of perturbation predicted by the model trained on gene knockout labels are also much stronger than those of the K562E-trained model across the board. Considering the 9.6% average training accuracy of the RPE1 gene model, a match probability of 19% is an excellent result, suggesting that the PPP6C gene may be experiencing direct perturbation in the OSD-91 samples, although this does not preclude the possibility that others are as well.

In terms of biological function, the top knockouts in RPE1 Cluster 0 are enriched for gene sets that are involved in the assembly of protein complexes, including those involved in RNA processing such as the Integrator Complex, as well as critical cell cycle functions, particularly cell membrane separation. They are also enriched for gene sets known to be responsible for physiological issues such as abnormal mitochondrial morphology and mental deterioration. The functional annotation of the second most probable cluster, RPE1 Cluster 24, is centered almost entirely around RNA processing and maturation, including capping and tail formation, methylation processes, and histone encoding. It also has a single overlap with a gene set involved in extra-nuclear transport. The complete gene set enrichment results for these clusters are in supplementary table 2.

The genes predicted to be knocked-out by the RPE1-trained model are consistent with the gene families identified as having DMHRs in the parent study for OSD-91. The gene PPP6C, for example, encodes a protein phosphatase subunit. While PPP6C itself was not identified in the original study, several other phosphorylation related kinase subunits, such as the PIK3CD are noted to have DHMRs. Similarly, the gene SLC1A5, which encodes solute carrier family 1 member 5 and is also a type of retrovirus receptor, is not reported directly, but SLC13A3, SLC16A5, SLC7A7, SLC7A8, SLC39A3, SLC23A2, and SLC29A1 are all reported as having DHMRs. GAB2, a specific binding protein for GRB2, is not reported, but GABRP is. Finally, for FAM136A, which is one of many genes grouped by familial sequence similarity, the most similar genes reported are FAM216A, FAM129B FAM184B, and FAM175B (Chowdhury et al., 2016). Only *MYBL2*, which is a MYB proto-oncogene, had no obvious presence or similarity in the reported gene lists (NCBI, 2024).

Overall, our pre-trained machine learning models identified genes and gene families in the OSD-91 simulated microgravity samples that were consistent with the findings of the original study, indicating that models pre-trained with Perturb-Seq data are able to identify reliable sources of genetic perturbation in new data.

## DISCUSSION

Transfer learning is a powerful technique in which knowledge gained while training a model in one domain, such as biology or medicine, can be applied to a related domain where there are fewer training data, such as rare diseases or space biology (Sanders, 2023). Transfer learning has been successfully applied to train classifiers to learn the underlying patterns in RNA sequencing data, with high accuracy prediction on small space biology datasets (Li, 2023). Here, we provide a substantial contribution to this field by training machine learning models to predict the origins of gene expression perturbations using single cell RNA sequencing datasets and the Perturb-Seq paradigm. We demonstrate the potential of Perturb-Seq data for training machine learning systems to predict gene expression behavior in complex experimental data. The increasing number of Perturb-Seq studies and the publication of GWPS mark a milestone in the transition of the method from a novel proposal to an established practice.

Here, we trained three different ML architectures to identify the sources of perturbation using the transcriptional profiles of individual cells, providing a comprehensive picture of the nuanced relationships of genetic cause and effect that propagate on a cell-wide level. We then used the best performing model to identify the sources of perturbation in the cells of a simulated spaceflight experiment, which were consistent with the findings of the original study. Our study successfully demonstrates the use of Perturb-Seq data for transfer learning to the space biology domain, compares multiple machine learning model architectures for this purpose, and provides insight to the origins of gene expression perturbation due to spaceflight stressor exposure. In particular, our study strengthens the previous findings that altered gravity exposure leads to perturbation of the expression of genes with DHMRs, but builds on these previous findings by identifying the specific genes likely responsible for downstream perturbation.

Our approach provides the unique ability to assign probabilities to individual genes as most likely to be perturbed to the extent that they are predicted to be completely “knocked out” by the experimental exposure. One interpretation of these probabilities is that they represent the level of perturbation a gene is experiencing. The ability to leverage this methodology to quantify the results of spaceflight exposures at the gene level lends a new facet of exploration to the precious datasets already generated by spaceflight experiments.

Future work may expand the utility of this approach in multiple ways. For example, our results identified the MLP architecture as the best performer, but using a system such as AutoKeras to converge on an optimum CNN configuration rather than manual hyperparameter tuning may yield a more robust NN architecture. Additionally, applying a form of One-Hot vector encoding to the data as an alternative method of normalization might increase its suitability for use in a CNN model, or computational methods such as ACTIONet, which incorporates archetypal analysis and manifold learning to offset cell cycle transcripts prone to scRNA-seq may also improve performance (Mohammadi et al., 2020).

Further, we provide recommendations for improving Perturb-Seq dataset generation and usage in future studies. 1) Datasets intended for practical application should be created in cell lines with consistent ploidy levels, thereby minimizing the gap between cultured cells and the tissue they represent as much as possible. 2) The coverage of the transcriptional profile should be as great a depth as possible. 3) Datasets should have a roughly equal distribution of sequenced cells for every knockout targeted. Additionally, in order to maximize the utility of existing Perturb-Seq datasets, it would be possible to expand upon the scoring method we presented here, to create a metric that accounts for knockout to cell count distributions, cluster to cluster composition contrasts, MOI, and depth of sequencing coverage in addition to the existing internal uniformity measure. This composite formula could then be applied universally to any Perturb-Seq dataset to provide a standardized quality score.

While the OSD-91 samples provide a sound metric for prediction performance, they represent a simulation of the biological effects of microgravity only. The next step in assessing the performance of the models is to make predictions on samples which have been to space, and ideally have been collected from humans instead of using immortal cell lines. The Inspiration 4 mission is a prime candidate for samples that meet these requirements (Jones, 2014). Additionally, as the models are trained on generalized perturbational data and can make predictions on any samples that are correctly formatted and express DEGs, their capacity to make predictions on a wide variety of samples should be explored in order to assess the versatility of the system as a universal tool.

For transfer learning, the importance of assessing and comparing multiple architectures in parallel becomes evident in light of the way our MLP model - which is a simplistic model algorithm by comparison - unexpectedly outperformed our CNN model in terms of accuracy without overfitting. While this outcome is both semantic and apt to change with further model development and optimization, it highlights how the method of pre-training models to learn generalizable knowledge serves as a primer for the more specific scope of space biology predictions is largely independent of algorithm type.

The ability to use large non-space biomedical datasets like Perturb-Seq for pre-training generalizable models to learn inherent biological interactions - in this case the responses of biological genes and pathways to specific gene knockouts and functional groups - is on its own an incredible insightful method of biological study. The ability to take the models trained on these datasets and use them to make predictions on unrelated small-volume datasets, especially those from spaceflight experiments, opens up an entire swath of analytical methods which would be otherwise precluded by data availability.

When refined, this system has the potential to provide a valuable tool for probing the seemingly inscrutable pattern of effects spaceflight has on biology.

## Data Availability Statement

The code written to create the models has been uploaded to Github, and the models themselves to Hugging face at the following locations: https://github.com/lauren-sanders/AI4LS https://huggingface.co/liamfj17/perturb_seq_prediction_models

All data used within the scope of this project is publicly available and can be found at the following locations:

Perturb-seq (2016) - https://www.ncbi.nlm.nih.gov/geo/query/acc.cgi?acc=GSE90063 Genome-Wide Perturb-Seq - https://gwps.wi.mit.edu/

or

https://plus.figshare.com/articles/dataset/_Mapping_information-rich_genotype-phenotype_landscapes_with_genome-scale_Perturb-seq_Replogle_et_al_2022_processed_Perturb-seq_datasets/20029387

OSD-91 Dataset - https://www.ncbi.nlm.nih.gov/geo/query/acc.cgi?acc=GSE65944

or

https://osdr.nasa.gov/bio/repo/data/studies/OSD-91

